# Imaging human cortical responses to intraneural microstimulation using magnetoencephalography

**DOI:** 10.1101/427815

**Authors:** George C. O’Neill, Roger H. Watkins, Rochelle Ackerley, Eleanor L. Barratt, Ayan Sengupta, Michael Asghar, Rosa Maria Sanchez Panchuelo, Matthew J. Brookes, Paul M. Glover, Johan Wessberg, Susan T. Francis

**Affiliations:** Sir Peter Mansfield Imaging Centre, School of Physics and Astronomy, University of Nottingham, Nottingham, UK; Department of Physiology, University of Gothenburg, Gothenburg, Sweden; Aix Marseille Univ, CNRS, LNSC (Laboratoire de Neurosciences Sensorielles et Cognitives – UMR 7260), Marseille, France

**Keywords:** magnetoencephalography, somatosensory cortex, intraneural microstimulation, single-unit

## Abstract

The sensation of touch in the glabrous skin of the human hand is conveyed by thousands of fast-conducting mechanoreceptive afferents, which can be categorised into four distinct types. The spiking properties of these afferents in the periphery in response to varied tactile stimuli are well-characterised, but relatively little is known about the spatiotemporal properties of the neural representations of these different receptor types in the human cortex. Here, we use the novel methodological combination of single-unit intraneural microstimulation (INMS) with magnetoencephalography (MEG) to localise cortical representations of individual touch afferents in humans, by measuring the extracranial magnetic fields from neural currents. We found that by assessing the modulation of the beta (13-30 Hz) rhythm during single-unit INMS, significant changes in oscillatory amplitude occur in the contralateral primary somatosensory cortex within and across a group of fast adapting type I mechanoreceptive 20 afferents, which corresponded well to the induced response from matched vibrotactile stimulation. Combining the spatiotemporal specificity of MEG with the selective single-unit stimulation of INMS enables the interrogation of the central representations of different aspects of tactile afferent signalling within the human cortices. The fundamental finding that single-unit INMS ERD responses are robust and consistent with natural somatosensory stimuli will permit us to more dynamically probe the central nervous system responses in humans, to address questions about the processing of touch from the different classes of mechanoreceptive afferents and the effects of varying the stimulus frequency and patterning.

## Introduction

In humans, our sense of touch is conveyed by an array of discrete low-threshold mechanoreceptive afferents 50 distributed throughout the skin, and haptic sensations are derived from a combination of activity across stimulated receptors (Johnson, 2001; Saal and Bensmaia, 2014). The sensory afferents from these receptors project to the somatosensory cortices, via the thalamus, enabling the decoding of the nature and location of the stimulus evoking the sensation. Our somatosensory abilities support a wide range of systems in the body, from integrating with the motor cortex for force feedback and fine-motor control (Ackerley and Kavounoudias, 2015), to emotional impact, providing feedback from social situations (McGlone et al., 2014).

Much of our knowledge about human mechanoreceptive afferents and their electrophysiological properties stems from microneurography studies (Vallbo and Hagbarth, 1968), where an electrode is placed in a peripheral nerve to register axonal impulses. Microneurography allows the characterisation of *single* mechanoreceptive afferent responses when the associated receptive field in the skin is excited by touch. Four distinct low-threshold mechanoreceptor classes exist in glabrous (non-hairy) skin, each with their own distinct spiking pattern in response to somatosensory stimuli and spatial representations within the skin (Vallbo et al., 1984a). In addition to registering impulses, it is possible to pass a small current (of approximately a few microamperes) down the electrode to elicit artificial impulses in the same mechanoreceptive afferent, in the absence of contact to the skin. Here, a small, corresponding simulated tactile sensation can be felt, i.e. the projected receptive field (at the same site as the physiological receptive field), which is thought to represent the sensation conveyed by these individual mechanoreceptors. This process has been termed single-unit intraneural microstimulation (INMS; Torebjörk and Ochoa, 1980; Vallbo, 1981) and allows precise impulse patterns to be evoked in individual nerve fibres, something traditional transdermal or intraneural stimulation cannot provide. The use of low current in INMS allows single afferents to be stimulated and avoids both the recruitment of numerous afferents and involuntary movements that may be induced by efferent activation when using higher currents, such as with median nerve stimulation. Single-unit INMS has shown that physiologically-defined mechanoreceptor types have distinct perceptual correlates in humans (Ochoa and Torebjörk, 1983; Vallbo et al., 1984b). Fast adapting type I (FA1) and type II (FA2) afferents elicit a vibration or fluttering sensation when stimulated, whilst slowly adapting (SA) type I (SA1) give rise to a pressure or pulling sensation. Slowly-adapting type II (SA2) afferents do not typically produce a clear, identifiable sensation. The type I units, which are more numerous (Johansson and Vallbo, 1979), provide small, point-like sensations. The use of single-unit INMS has the advantage over traditional tactile stimuli of being able to produce a defined and controllable pattern of firing within a single afferent and single afferent class. This allows the precise examination of how input patterns from peripheral mechanoreceptive afferents are perceived and represented in the central nervous system.

Despite knowledge of the peripheral and perceptive properties of single mechanoreceptive afferents, little is known about how their responses to stimulation manifest in the human brain. Using single-unit INMS, it is possible to probe the sense of touch at the “quantal” level of individual afferents, something which cannot be achieved for any of the other senses. There have been relatively few studies in which neuroimaging has been combined with single-unit INMS to investigate the neural effects of selective mechanoreceptive afferent stimulation. A study by Kelly et al., (1997) used high density electroencephalography (EEG) over the primary somatosensory cortex (S1) and showed that stimulating single FA1 or SA1 units projecting to the hand gave characteristic peaks in power at the same frequency at which the units were stimulated. Only two studies have been performed using functional magnetic resonance imaging (fMRI) to map the spatial profile of the cortical responses to single-unit INMS (Sanchez Panchuelo et al., 2016; Trulsson et al., 2001). The earlier study, performed at 3.0 T, showed that individual type I units have corresponding neural activations in S1 and the secondary somatosensory cortex (S2), and the responses were in good spatial agreement with activations from vibrotactile stimulation (Trulsson et al., 2001). The more recent study was performed at ultra-high field (7.0 T) using high spatial resolution to show that functional activation patterns of units located on the hand digits were spatially well-localised within the expected digit regions, ascertained from maps of digit somatotopy obtained from vibrotactile stimulation to the skin of the fingertips (Sanchez Panchuelo et al., 2016).

Magnetoencephalography (MEG) measures the extracranial magnetic field associated with extracellular current flow. MEG provides unique spatiotemporal resolution, offering vastly improved temporal resolution over fMRI (which is limited by the haemodynamic lag of the BOLD effect, and it being an indirect measure of neural activity) and an improved spatial resolution compared to EEG (as magnetic fields are not distorted by the skull to the same extent as electric potentials). In this paper, we performed single-unit INMS, using a recently developed system designed for compatibility with MEG (Glover et al., 2017). We aimed to demonstrate that this approach can localise the cortical response to single-unit INMS and capture the spatiotemporal dynamics of responses to simulation of individual mechanoreceptors, providing the feasibility of performing future studies to probe how the dynamics of different patterns of activity in these mechanoreceptors are represented cortically.

## Methods

### Experimental Procedure

A total of 9 healthy participants (age 24-68 years, 5 female) volunteered to undergo microneurography/single-unit INMS and simultaneous MEG recording. Each participant was given detailed information about the procedure and provided their written informed consent. Ethical approval was granted by the University of Nottingham Medical School Ethics Committee, with all procedures conducted in line with the Declaration of Helsinki.

Experimental sessions consisted of 3 phases: 1) microneurography for the characterisation of a single mechanoreceptive afferent (Vallbo and Hagbarth, 1968); 2) assessment of the sensation to single-unit INMS; 3) MEG recordings during INMS. These steps were repeated and ∼2-3 individual mechanoreceptive afferents were typically stimulated during MEG data collection in a single experimental session. The participant was seated comfortably with their head just below the MEG helmet, and were instructed to sit as still as possible with their arm/hand held still using a vacuum cushion to minimise any slight movement of the arm/hand, which would result in slight movements of the electrode and loss of the unit. A high-impedance, insulated tungsten recording/stimulating electrode (15 mm length, shaft diameter 200 µm, tip diameter ∼5 µm; FHC, Bowdoin, ME) was then inserted percutaneously into the left median nerve, approximately 30 mm proximal from the wrist crease. A second uninsulated reference electrode was placed subcutaneously 50 mm away from the first and both were connected to an INMS system specifically designed to operate without generating measurable magnetic interference (Glover et al., 2017). A diagram of the electrode setup is shown in Figure 1.

**Figure 1.**
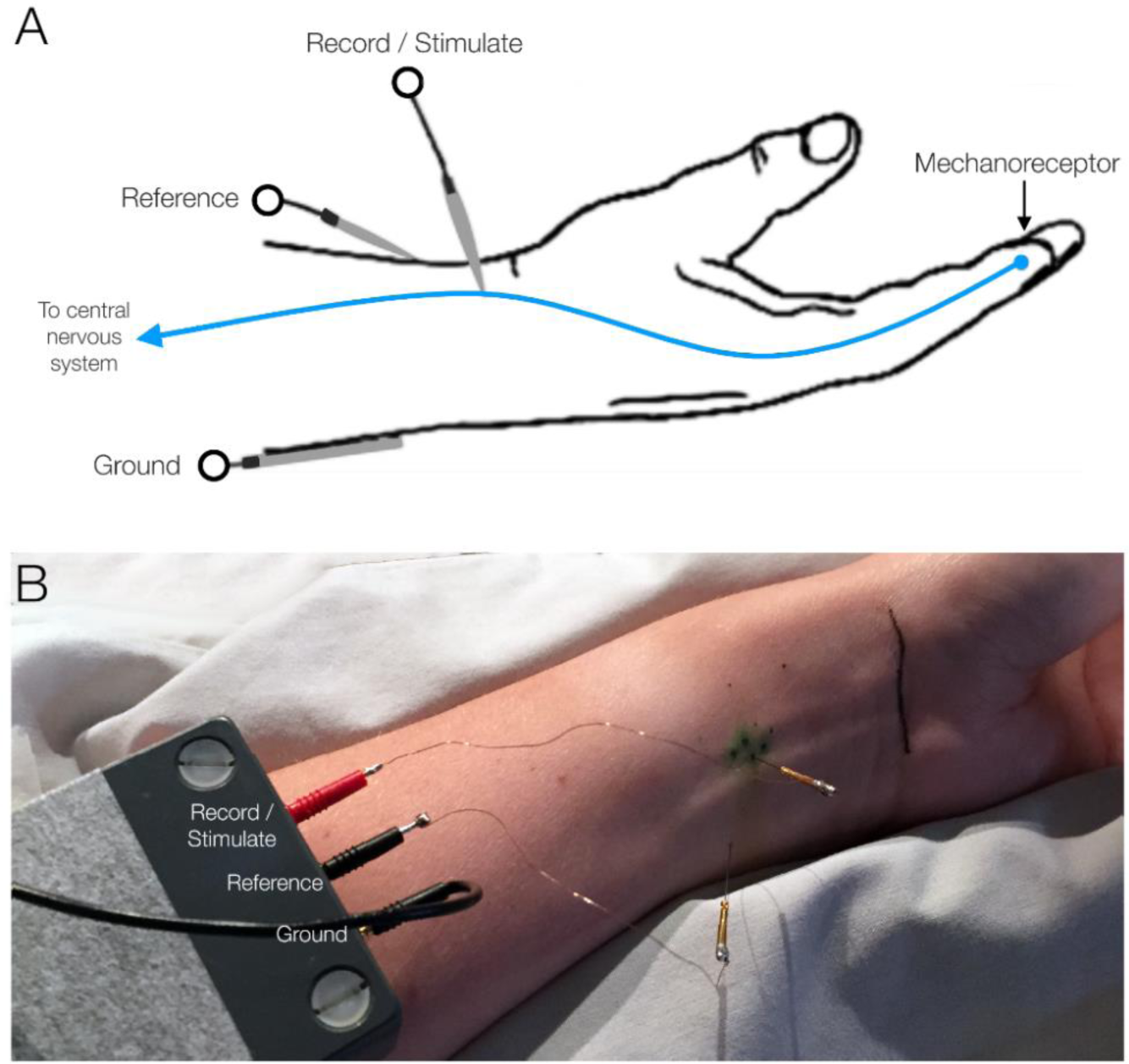
Experimental setup for microneurography and single-unit microsimulation. A) A simplified diagram showing how a single mechanoreceptive afferent is interrogated. A recording/stimulating electrode is placed in the median nerve to register activity from a single nerve fibre. If mechanical stimulation is applied to the afferent’s receptive field, the electrode in ‘record mode’ will register the spiking pattern being sent to the central nervous system. In ‘stimulate mode’, passing a small current down the electrode will stimulate the mechanoreceptive afferent, eliciting a tactile sensation at the location of the receptive field in the skin. B) The experimental setup on a participant in the study.

For experimental phase 1, the INMS system was in ‘record mode’, and a real-time visual and audio readout of the potential difference between the two electrodes was output to the experimenter within the magnetically shielded room. The location of mechanoreceptive afferents was determined by gentle touch across the hand, delivered by the experimenter, between minor adjustments of electrode position, until a unitary recording was detected. The precise location of the unit receptive field on the skin and force activation threshold of the unit was then identified using von Frey monofilaments and marked with ink. Once the mechanoreceptive afferent was identified, the response pattern and spatial extent on the skin was examined to identify the receptor type as SA1, FA1, SA2 or FA2 (Vallbo et al., 1984a). The INMS system was then switched to ‘stimulate mode’, to assess the sensation to INMS (experimental phase 2). A train of constant current pulses (pulse width 200 µs, amplitude <10µA) was delivered at 60 Hz for 1 s. The trains of pulses were delivered repetitively and the current was increased gradually (resolution 0.1 µA), until the participant felt a sensation in their hand, which was time locked to the stimulation. The participant was asked to describe the nature and location of the sensation; if there was an exact correspondence with the physiological properties of the unit (e.g. location, quality, size) with only a single unit stimulated at that current, and the investigation proceeded to the MEG data collection (experimental phase 3). The paradigm during the MEG recording consisted of 80 trials of a 1 s long train of pulses (as described above), with a base inter-trial-interval of 10± 1 s to jitter the stimulus. After every 10 trials, the quality and intensity of the sensation was confirmed verbally with the participant, as small movements of the electrode can diminish the intensity of the sensation, presumably reflecting failure to generate impulses with each stimulation pulse. In the case where a participant no longer felt the sensation, the pulse amplitude was increased until a similar quality of sensation was again felt again in the same location (Torebjörk et al., 1987). If a comparable sensation was achieved by increasing the stimulation intensity, the MEG protocol was resumed, but if comparable sensation intensity was not recovered, or there was evidence of a change in the unit stimulated (e.g. its quality/location), the MEG recording was terminated.

Immediately after INMS, the participant underwent a short experiment, where mechanical vibrotactile stimulation was applied to the skin at each microstimulated unit’s receptive field location to determine the amplitude at which the vibrotactile stimulation matched the INMS sensation. A small piezoelectric vibrotactile stimulator (Dancer Design, St. Helens, UK) controlled by an Arduino microcontroller (http://arduino.cc) was used to deliver a vibrotactile stimulation at 60 Hz. The amplitude was adjusted until the sensation felt similar in intensity to that from the INMS. These amplitudes were noted and used in a subsequent MEG experiment in which vibrotactile stimulation at each microstimulated unit’s receptive field was carried out with identical timings to the INMS paradigm. This vibrotactile/MEG follow-up session was performed at a later date to allow for the maximum number of units to be stimulated within each INMS experimental session.

### Data Acquisition

MEG data were recorded using a 275 channel MEG system (CTF; Coquitlam, BC) in synthetic 3^rd^ order gradiometer configuration, with a sampling rate of 1200 Hz, and a hardware anti-aliasing low-pass filter of 300 Hz applied. An analogue trace of the INMS stimulation pulses was recorded concurrently to allow for offline synchronisation of the neuromagnetic recordings. All participants wore a bespoke flexible headcast (details of the manufacturing procedure can be found in Liuzzi et al., 2017; Meyer et al., 2017), designed to minimise head movement within the MEG system. The headcasts include cavities designed to fit three head position indicator (HPI) coils, which were periodically energised during the experiment, to determine their position within the MEG sensor array. The HPI coil locations relative to the brain were defined during headcast manufacture, meaning 170 potential source localisation errors typically introduced when misaligning the sensors and the anatomy by other means were minimised (Troebinger et al., 2014). For a single participant, the headcast failed to fit and so an alternative method to coregister the brain to the sensors was employed. In this case the HPI indicators were attached to the head at the nasion and bilateral preauricular locations. A measurement of coil locations relative to the scalp was recorded using a 3D digitiser (Polhemus; Colchester, VT). Coregistration was achieved by matching the digitised surface with a surface extracted for the anatomical image. Anatomical images were acquired using a 3.0 T MR System (Philips Healthcare; Best, Netherlands) using an MPRAGE sequence (1 mm^3^ isotropic resolution).

### Data Analysis

MEG data were visually inspected for artefacts. If an epoch contained excessive artefacts (for example generated by muscle or eye movement, eye blinks, or from SQUID resets), it was omitted from further analysis. The sensor data were then band-pass filtered to the beta band (13-30 Hz), as this is one of the oscillatory components modulated by somatosensory stimulation (Hari and Salmelin, 1997; Cheyne, 2013). MR images were segmented and cortical surfaces were reconstructed using FreeSurfer 5.3 (Dale et al., 1999; Fischl et al., 1999). Further segmentation of the scalp, as well as inner and outer skull surfaces, were generated using modified routines from FieldTrip (Oostenveld et al., 2011). The surface which defined the grey/white matter boundary was decimated to 12500 vertices per hemisphere and each vertex was used as a location for source reconstruction. Source reconstruction at each vertex was achieved with a Linearly Constrained Minimum Variance (LCMV) beamformer, with the data covariance matrix generated from the entire duration of the experiment (Brookes et al., 2008). Dipole approximation for forward modelling was calculated using a 3-shell boundary element model (BEM; Stenroos and Nummenmaa, 2016), with the optimal dipole orientation for maximal signal variance derived using an eigenvalue decomposition approach (Sekihara et al., 2004). With beamformer weights for a given vertex, *w*, generated, they are employed to quantify changes in power across the cortex. For every location, an activation index, *A*, was generated to quantify changes in source power during the stimulation (0-1 s after stimulus onset) compared to a baseline period (8.9-9.9 s after stimulus onset). For the *i*^*th*^ trial, *A* was calculated at each location using Equation 1,

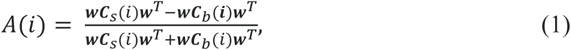

where *C*_*s*_(*i*) and *C*_*b*_(*i*) are the single-trial sensor-level covariance matrices which correspond to the stimulation and baseline epochs respectively. Activation indices were averaged across trials to generate a final activation image for each experiment. The activation index is scaled between m−1 and 1, with 0 denoting no modulation in power between the two epochs. To allow for a group level analysis, all images were transformed to a standard cortical surface derived from the MNI-305 template brain. In addition, to interpret source localizations multimodal parcellations published in the Glasser atlas (Glasser et al., 2016) were used. The Glasser atlas is a multi-modal parcellation of the brain derived from Human Connectome Project (HCP-MMP1.0) and was converted using the HCP Workbench software to fit the MNI-305 brain. In addition, we also compared our results to previously derived functional markers. Here we used probabilistic atlas of the hand area generated from fMRI responses to vibrotactile somatosensory stimulation of the left hand (Sengupta et al., 2018).

### Statistical Inference of Significant Power Changes

To identify which locations corresponded to a significant change in power during stimulation (INMS or vibrotactile), we employed non-parametric permutation testing (Maris and Oostenveld, 2007; Nichols and Holmes, 2002; Pantazis et al., 2005; Singh et al., 2003) at both the individual subject and group levels. At the individual subject level, our null hypothesis was that the oscillatory power measured in the stimulation and the baseline epochs within a trial were identical. Therefore, if we were to randomly shuffle the labelling of the covariance matrices within trials, prior to generating *A* in Equation 1, then averaging across trials to form a null averaged activation index, Ã, would not give a significantly different result to the genuine averaged activation index, Ā. 2000 permutations of Ã were generated to create a null distribution for each vertex location. To correct for multiple comparisons, the threshold free cluster enhancement score (TFCE; Smith and Nichols, 2009) was calculated for every vertex on each iteration of Ã. For a given location *r*, the TFCE score was calculated as

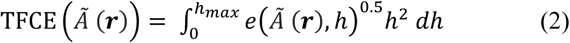

where *e*(*Ã* (*r*)*, h*) is a function which returns the spatial extent of a cluster at *r*, should *Ã* (*r*) exceed a threshold value *h*, and 0 otherwise. Positive and negative clusters were treated as non-contiguous, and the maximum absolute value of TFCE on each iteration was used to generate a global null. Any vertex where the absolute TFCE of *Ā* exceeded the 95^th^ percentile of this null was considered significant.

At the group level, our null hypothesis was that the activation maps simply represented zero-mean noise, so randomly multiplying any of the experiments’ trial averaged activation images by −1 and then group averaging to generate a null activation map, *Ã*, would not give a significantly different result to the group-averaged image *Ā*. All *n*! possible permutations were calculated and similarly to the group level analysis, multiple comparisons correction was handled by finding the critical TFCE threshold. By performing statistical analyses at both the group and individual level, we can determine whether changes in oscillatory power which are observed at the group level changes are potentially a) applicable markers for studying changes in individual experiments and b) generate a probabilistic map of where the cortical source of stimulation to the hand area localises to.

## Results

### Electrophysiology

Across the cohort of participants, 39 single-unit recordings were identified. 19 of these provided percepts that matched the unit physiology, indicating successful single-unit INMS (Torebjörk et al., 1987) and cortical responses to stimulation were recorded during MEG. An excerpt from a single-unit recording from an FA1 can be seen in Figure 2A. Here, we observed high spiking rates (occasionally exceeding 200 impulses.s^−1^) when a von Frey monofilament was pressed into the receptive field location in the skin. Data from 3 units were discarded due to too few MEG trials being recorded before losing the unit sensation. The properties of each of the 16 units with full MEG recording protocols can be found in Table 1, and their locations on the hand in Figure 2B. 10 units were identified as mechanoreceptor type FA1, 5 as SA1, and 1 as FA2. In the results presented here, we primarily focused on the 10 FA1 units, given that these formed the largest group, and their perceived sensation — vibration/flutter — is similar to that of vibrotactile stimulation. For completeness, the localisation results of the other types, 5SA1 units and a single FA2 unit, can be found in the Supplementary material.

**Table 1.** Metadata on the 16 microstimulated units recorded in the MEG scanner. Unit identifiers are based on the order of collection. Note all locations are on the palmar side of the hand unless otherwise noted (D denotes digit number).

| Participant ID | Unit ID | Type | Location | Sensation | Threshold current (uA) | Perceptive Field Size (mm) |
| --- | --- | --- | --- | --- | --- | --- |
| 1 | 1a | FA1 | Medial D3 | Vibration | 2.2 | Not recorded |
|  | 1b | SA1 | Proximal D4 | Pulling | 0.9 | 0.1 |
| 2 | 2a* | FA1 | Distal D3 | Tingle | 2.2 | 2 |
| 3 | 3a | SA1 | Palm | Pulling | 1.4 | 0.1 |
|  | 3b | SA1 | Palm | Pulling | 2.8 | 1 |
|  | 3c | FA1 | Palm | Vibration | 3.7 | 1 |
|  | 3d | FA1 | Distal D3 | Flutter | 3.3 | 1 |
|  | 3e | FA1 | Dorsal Medial D3 | Flutter | 0.9 | 1 |
| 4 | 4a | FA1 | Distal D4 | Vibration | 3.4 | 1 |
|  | 4b | FA1 | Distal D4 | Vibration | 4.2 | 2 |
|  | 4c | FA2 | Distal D4 | Tapping | 4.4 | 7 |
|  | 4d | SA1 | Medial D4 | Point-like pressure | 2.4 | 0.1 |
|  | 4e | SA1 | Medial D4 | Pressure | 3.2 | 3 |
|  | 4f | FA1 | Distal D4 | Vibration | 2.0 | 3 |
| 5 | 5a | FA1 | Distal D1 | Tingle | 4.2 | 2 |
| 6 | 6a | FA1 | Palm | Prickle | 1.1 | 3 |

\*Unit acquired without headcast

**Figure 2.**
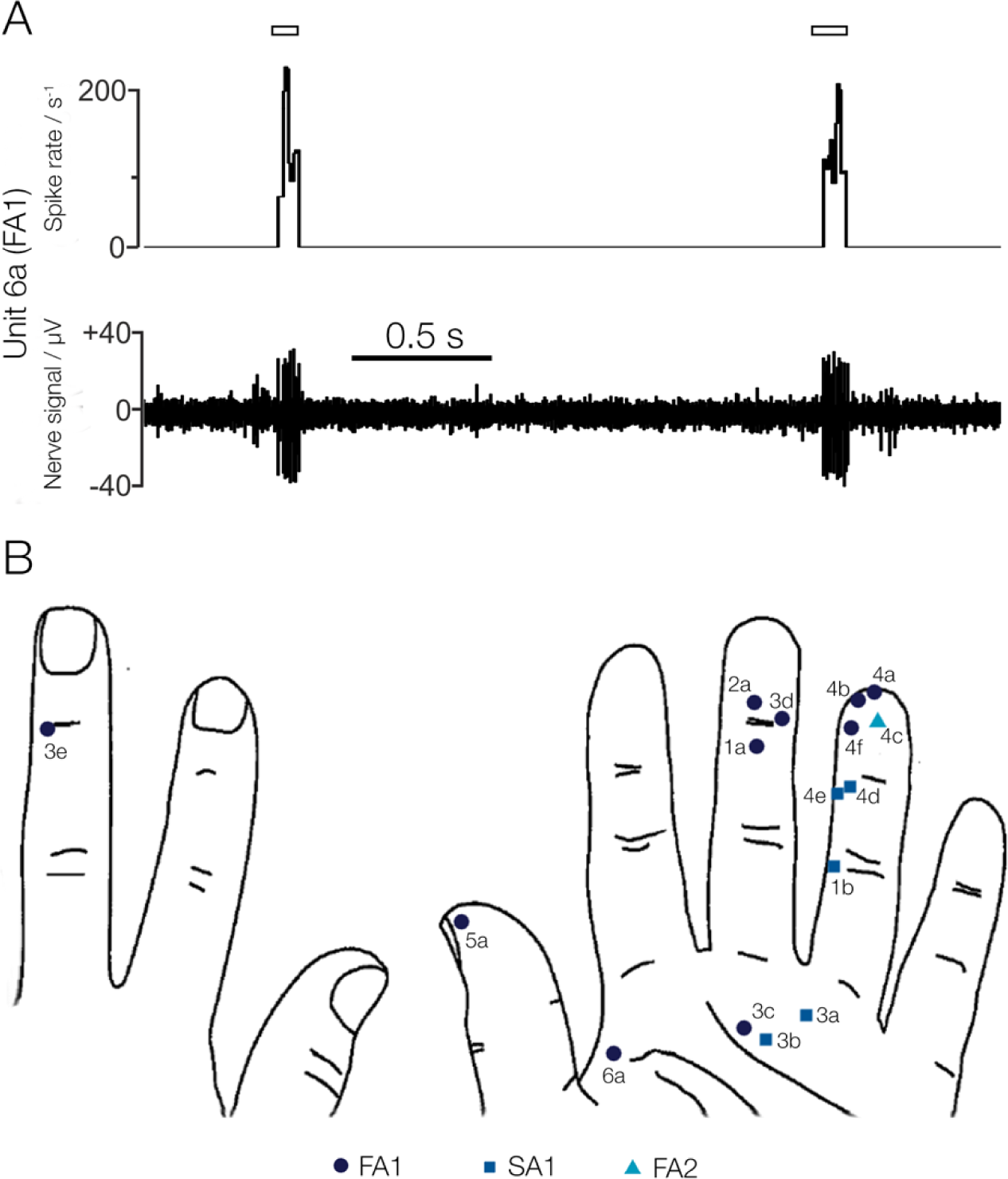
A) Excerpt from a single-unit peripheral neurve recording from a fast adapting type 1 (FA1) afferent (unit 6a), showing the unitary responses to mechanical stimulation of the receptive field in the skin whilst in record mode. Upon single-unit INMS, the subject felt a 3 mm spot defined to have a ‘prickle’ sensation. The location of the receptive field can be found in panel B. (B) Receptive field locations in all 16 units in the right hand which were microstimulated whilst in the MEG scanner, with adjacent numbers corresponding to the Unit ID in Table 1. The shapes correspond to the receptor type; dark blue circles represent FA1 afferents (vibration sensation in a very small area); blue squares show SA1 afferents (pressure or pulling in a small area), and the cyan triangle represents an FA2 afferent (diffuse vibration sensation).

### MEG

Figure 3 identifies where in the brain beta power changes during single unit INMS across the 10 FA1 units were 280 located. Figure 3A shows a conjunction map of the number of individual units that had a significant (*p*_*corrected*_ < 0.05) change in power. We observed a large degree of overlap across the units in the contralateral primary somatosensory and motor (M1) cortices, with 8/10 units in spatial agreement; the other 2 units showed no significant changes in power at any spatial location in the brain (see Supplementary material). In the ipsilateral S1, we observed overlap of activations in 2/10 units. Figure 3B shows the map of group-averaged activation index (*Ā*_INMS_), thresholded to show only the significant changes in beta power in response to INMS at the group level. This map shows that there was a strong decrease in oscillatory power, which was (much like the high overlap region at the individual level) situated primarily over the contralateral S1 and extended into M1. However, it is important to note that these three areas with the largest reductions in beta power could be in fact a single diffuse reconstructed source that is represented in multiple locations when constrained to a cortical surface. To help clarify where the origin of this source may be, the location of maximal reduction is represented as a magenta dot and is situated in S1. Next, we compared our results to previous findings; Figure 3C is a zoomed representation of the contralateral sensorimotor area, with the Brodmann areas from the Glasser atlas (Glasser et al., 2016) and fMRI derived hand area (Sengupta et al., 2018) in grey and black, respectively. Here, we see that the area of with the largest spatial overlap (yellow) is primarily situated within the hand area. The magenta dot from Figure 3B has also been overlaid and is situated within the high spatial overlap area, implying the largest reduction in beta power is a common feature across units, rather than being driven by a small subset. Figure 3D is a zoomed representation of the findings in Figure 3B, Again, the Glasser atlas is overlaid, but in green for better contrast, and the hand area remains in black. The location of the peak reduction in beta power, represented by the magenta dot, is situated on the borders of Brodmann areas 1 and 2, confirming that the peak modulation of power occurred in S1, rather than in M1. Given that the magenta marker was situated at the largest group effect, and the highest amount of overlap across individual subjects, this makes for a reasonable candidate location to place a virtual electrode to explore the other spectral and temporal properties during the experiment. The results of such are shown in Figure 3E, which shows a group-averaged time-frequency spectrogram of INMS at this location. The spectrogram shows that it is the beta oscillatory band that showed the largest fractional change in power during stimulation, where we observed event-related desynchronisation (ERD). After the stimulation ceased (t = 1), post-event rebound (PER) occurred, with an overshoot in beta power at t = 2-3 s, before returning to baseline. Performing the same localisation analysis on beta power in the PER epoch showed this to be significant at the group level, but individual testing showed that this was only significant for three units, all from the same participant (see Supplementary material). We also found similar temporal properties in the t = 0-1 s and t = 2-3 s epochs in the alpha band, but few units showed individual level significance and the locations of group level significance had a reduced activation index magnitude compared to beta band activity (see Supplementary material).

**Figure 3.**
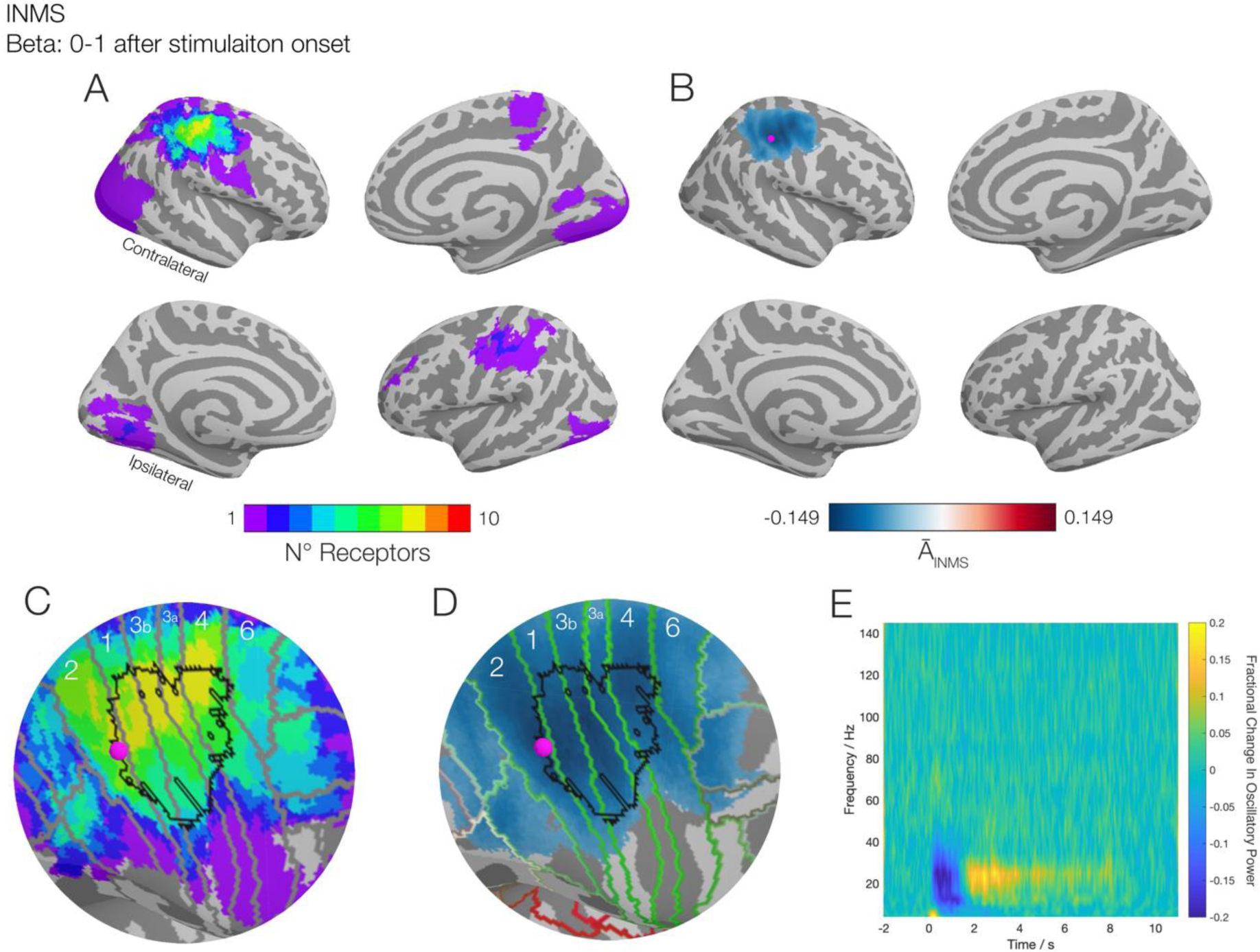
Results of the INMS from the group of 10 FA1 units studied overlaid onto inflated cortical surfaces, gyri are represented as light grey surfaces sulci are dark grey. A) Results of the individual unit analysis in beta band (13-30 Hz) power for a given location, represented as a conjunction map, showing the number of FA1 units that exhibit significant changes. B) Group-averaged activation image of the 10 FA1 units at a statistical threshold (p < 0.05), showing microsimulation elicits a reduction in beta oscillatory power over the contralateral sensorimotor area, with the peak reduction in beta power represented by the magenta dot) occurring in the postcentral gyrus. C) Zoomed portion of panel A, with the Brodmann areas as defined by the Glasser atlas (Glasser et al., 2016) overlaid in grey and hand area derived from a functional atlas (Sengupta et al., 2018) in black. Here the region representing the largest overlap across units (yellow; N=8) is contained primarily in the poscentral gyrus and within the predefined hand area. D) Zoomed portion of panel B, with both the Glasser atlas (this time in green) and the hand area overlaid. The peak reduction of beta power represented as the magenta dot) is located in postcentral gyrus. E) A time-frequency spectrogram of the peak location from panels B, C and D, showing a distinct event-related behaviour in the beta band which extends beyond the initial 1 s of stimulation. Interactive versions of the cortical plots can be found at http://georgeoneill.github.io

The equivalent statistical maps from the vibrotactile experiments are shown in Figure 4. In Figure 4A, we observed similar results at the individual level, as depicted in Figure 3A, with spatial overlap between the vibrotactile stimuli applied at the receptive field location for each unit and INMS in contralateral S1 in 9/10 corresponding vibrotactile stimuli. In the ipsilateral hemisphere we saw a greater number of overlapping responses between vibrotactile stimulation and INMS (3/10) with the area of overlap more parietal than contralateral S1. Figure 4B shows the group-averaged activation index (*Ā*_Vibrotactile_), again with a statistical threshold. A significant decrease in beta band oscillatory power was found over the contralateral postcentral gyrus during vibrotactile stimulation, agreeing with the findings of INMS. Furthermore, the magenta dot, which represents the location of the maximum reduction of beta power during stimulation, was located in the exact same vertex as the INMS experiments (i.e.0.0 mm distance between INMS and vibrotactile peaks). Figure 4C is a zoomed-in representation of the plot in Figure 4A, which again overlays the Glasser Atlas and the fMRI derived hand area. We observed that our regions exhibiting the highest level of significance across the subjects occurred within the hand area, and within the Brodmann Areas that corresponded to the postcentral gyrus. Figure 4D is a zoomed-in version of the group level image. We see that the peak location also occurred within the hand area. We note that this location coincided with one of the regions where N=9 in Figures 4A and 4C. The time-frequency vibrotactile data is shown in Figure 4E, where we observed a similar behaviour in the beta band, whilst the alpha band showed a rebound starting later than for the beta band, which may be related to slight movement of the finger in response the vibrotactile stimulation (Caetano et al., 2007), a finding not observed with INMS. Analyses for the alpha band during the stimulation and resynchronisation epochs, and the beta in the resynchronisation epoch, can be found in the Supplementary material, but none of these other conditions showed a high likelihood of significance in individual subjects.

**Figure 4.**
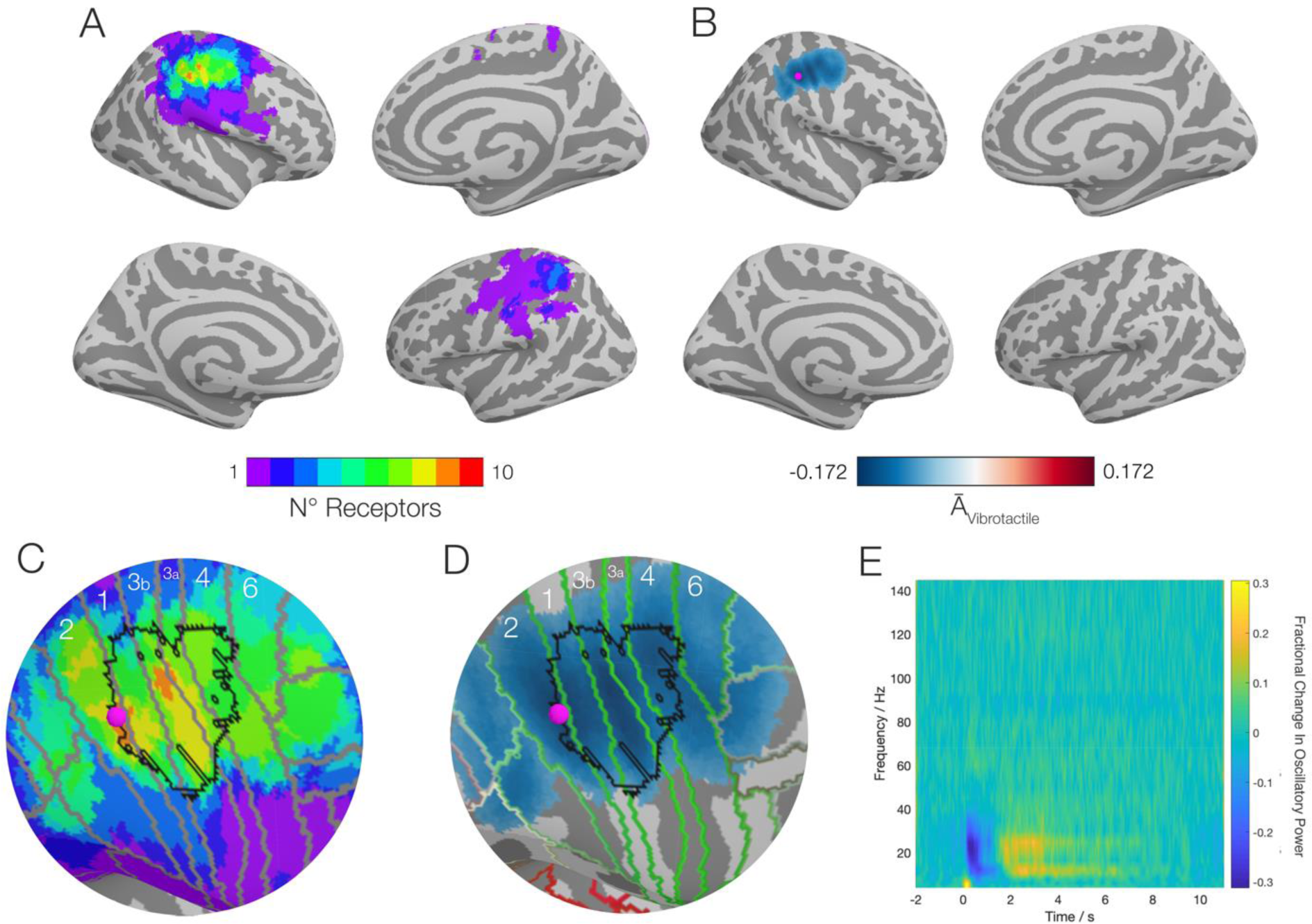
Results of vibrotactile stimulation at the perceived receptive field of the 10 FA1 units studied overlaid onto inflated cortical surfaces, gyri are represented as light grey surfaces sulci are dark grey. A) Results of the individual vibrotactile analysis, represented as a conjunction map, showing how many exhibited significant changes in beta band (13-30 Hz) power for a given location. B) The group-averaged activation image of the 10 vibrotactile images, with a statistical (p < 0.05) threshold applied, showing vibrotaction elicited a reduction in beta oscillatory power over the contralateral sensorimotor area. The location of the maximal reduction of beta power is represented with the magenta dot. C) Zoomed portion of panel 340 A, with the Brodmann areas as defined by the Glasser atlas (Glasser et al., 2016) overlaid in grey and hand area derived from a functional atlas (Sengupta et al., 2018) in black. Here the region representing the largest overlap across units (orange; N=9) is contained primarily in the Brodmann areas 2 and 3b, both S1 areas. D) A zoomed in portion of panel B, again with Glasser atlas (green) and hand area (black) overlaid. The peak reduction of beta power (represented as the magenta dot) is located in postcentral gyrus. E) Time-frequency spectrogram of the peak location from panel B, showing a distinct event-related behaviour in the beta band which extends beyond the initial 1 s of stimulation. Interactive versions of the cortical plots can be found at http://georgeoneill.github.io

Figure 5 shows the spatiotemporal similarities of responses to single-unit INMS and vibrotactile stimulation. Figure 5A provides a conjunction map representing the number of unit locations for which there was a significant change in beta power in both the INMS *and* corresponding vibrotactile experiments. Here, we saw that for any 350 given location, there was a maximum overlap in 8 subjects over the contralateral S1, whilst in the ipsilateral hemisphere only activation in a single subject showed co-significance. Further, at the group level, permutation testing found no significant differences between *Ā*_INMS_and *Ā*_Vibrotactile_ in the beta band during stimulation at any location within the brain. Figure 5B shows the conjunction map within the right S1 and M1. The map shows three features; locations where *Ā*_INMS_ was within 90% of the most extreme value (magenta), *Ā*_Vibrotactile_ within 90% of the most extreme value (cyan), and areas from Figure 5A where *n* = 8 (yellow). Overlaying these highlights two regions in S1 where all three conditions were satisfied (marked in black), one within Brodmann Area 3b, and one which straddles the borders of Brodmann areas 2 and 1, the latter of which agreed with the peak locations of *Ā*_INMS_ and *Ā*_Vibrotactile._ However, it is important to note that due to the nature of unfolding source reconstructed data onto a cortical surface, that these two distinct regions may be potentially the same source projected onto both sides of the gyrus. Figure 5C shows the group-averaged oscillatory power in only the beta band and compares the INMS beta timecourse to that of the vibrotactile experiments in the same location. The shaded areas represent the standard error across subjects. Non-parametric randomisation testing of the timecourses revealed no significant differences at any time point between the INMS and vibrotactile experiments.

**Figure 5.**
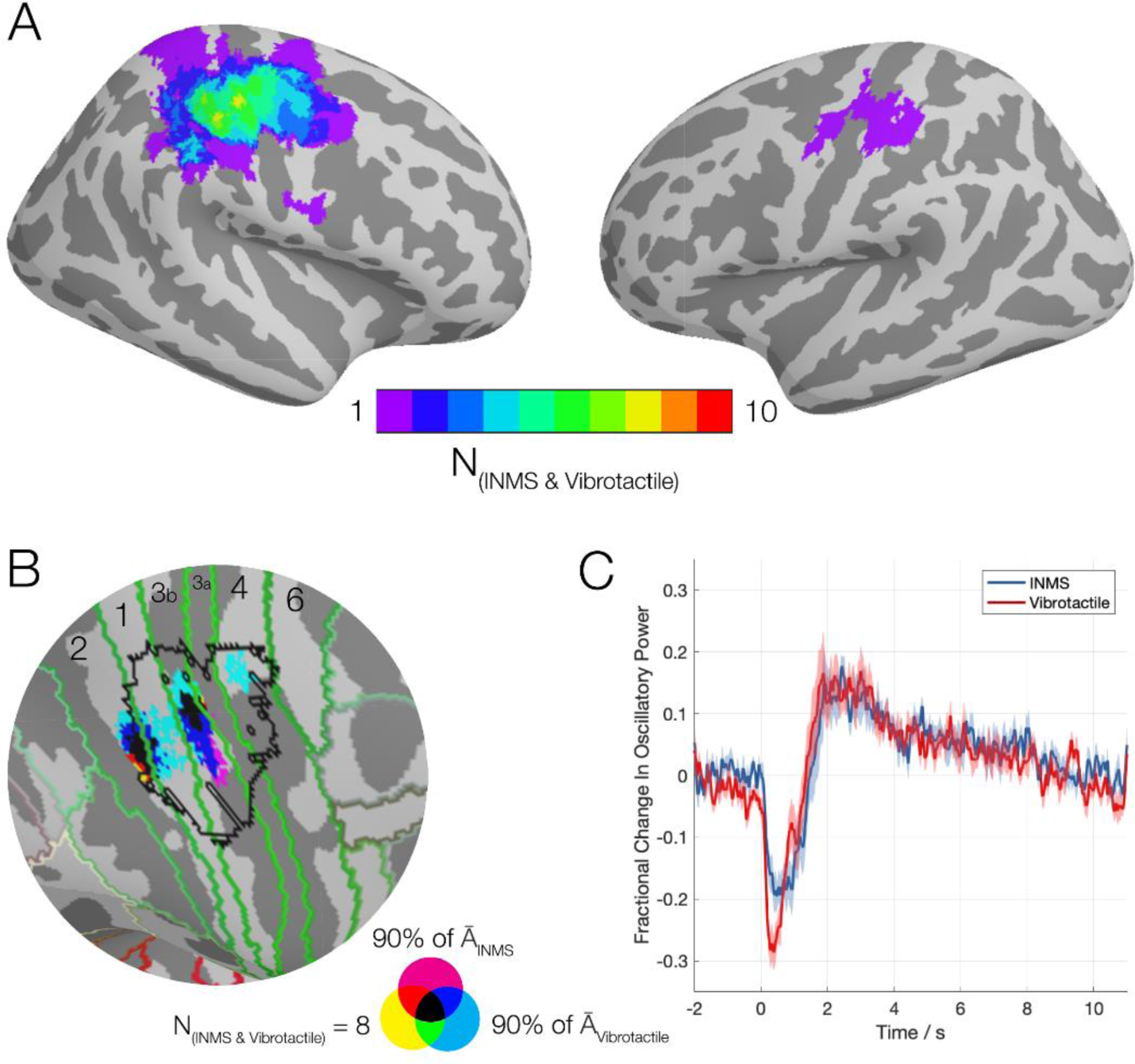
Comparison of the spatiotemporal properties between INMS and the vibrotactile experiments. A) A conjunction map of where both INMS and vibrotactile experiments showed significant changes in beta power for a given mechanoreceptive afferent, and how many experiments spatially overlapped for a given area. B) A zoomed map of the right hemisphere’s central gyri showing the largest group-averaged changes in power for INMS (magenta) and vibrotactile (cyan) and where they overlap with the locations of the largest spatial agreement as depicted in panel A (yellow). Any location where all three conditions held are represented in black. Brodmann areas, as defined using the Glasser atlas, and a fMRI derived hand area have been overlaid for reference. C) The group-averaged beta power timecourses for INMS (blue) and vibrotactile (red) responses, showing similar response timecourses in the contralateral (right) S1.

## Discussion

Here, we aimed to show the feasibility of combining single-unit INMS (using recently-developed stimulation hardware; Glover et al., 2017) with MEG, to probe whether we could reliably capture the spatial, spectral, and temporal properties of the cortical projections of individual single-unit mechanoreceptive afferents in human glabrous hand skin.

Our results show that single-unit INMS of these quantal elements of touch yielded a robust reduction of oscillatory power in the beta (13-30 Hz) band, which could be localised spatially using MEG, with 80% of individual FA1 units showing significant oscillatory power reduction in contralateral S1. Furthermore, we compared our results with oscillatory responses to amplitude-matched vibrotactile stimulation of the same receptive field location and showed that our ability to localise the ERD of individual mechanoreceptive afferent stimulation using INMS was as reliable as localising sources derived from vibrotactile stimulation and that the group results from single FA1 mechanoreceptive afferent stimulation revealed similar spatiotemporal patterns to those from vibrotactile stimuli. In addition, we also observed beta ERD to SA1 and FA2 units (see Supplementary material), showing that this was not a specific feature to FA1 units. The fundamental finding that single-unit INMS ERD responses were robust and consistent with natural somatosensory stimuli permits us to more dynamically probe central nervous system responses in humans. This will allow us to address a number of questions about the processing of touch from the different classes of mechanoreceptive afferent to a range of input patterns, including how varying the 390 stimulus frequency and patterning in the different receptor types may modify cortical responses.

In the current study we chose to assess beta power, which was motivated by a number of studies observing modulations in beta power to tactile stimulation using both intracranial (Brovelli et al., 2004; Crone et al., 1998; Witham and Baker, 2007) and extracranial modalities (examples include, but are not limited to Bauer et al., 2006; Cheyne et al., 2003; Gaetz and Cheyne, 2006; Pfurtscheller et al., 2002; Salenius et al., 1997; van Ede et al., 2010). Bauer and colleagues used a spatial-selective tactile task to investigate how attention affected oscillatory power during tactile stimulation. They found the reduction of beta power during stimulation was not significantly affected by attentional modulation, suggesting this was likely an automatic response to afferent stimulation, independent of processing demand (Bauer et al., 2006). The same study also noted that the amount of post-stimulation resynchronisation in the somatosensory cortex was reduced if participants were instructed to attend to the stimulus, compared to those who were asked to ignore. The sensation from microsimulation is subtle and required subjects to attend to it, thus we expected a high level of attention. It is thus interesting to note that stimulation in only a few of the units produced significant rebounds of beta power, and these all originated from the same participant, a participant well accustomed to single-unit INMS and so more familiar to the sensation, thus potentially requiring less attentional load. Another study has shown that anticipation may modulate event-related reductions in beta, in research employing a delay between a cue and the delivery of a stimulus, where there was contralateral ERD of the beta band, with the extent of the reduction monotonically increasing with delay time, implying an anticipatory element of somatosensory beta (van Ede et al., 2011). However, in the present study we used no explicit cues and therefore have confidence that our observations in beta power reduction are entirely due to stimulation.

Recently, there has been an emerging shift in how we interpret beta band oscillations in the somatosensory and motor cortices (van Ede et al., 2018). Rather than treating beta as a sustained oscillation, which is power modulated based on processing demand, it may be better to discuss beta in terms of transient (∼50 ms) events, or bursts. These punctate events have been shown to be robust across multiple species (Sherman et al., 2016; Shin et al., 2017), and their occurrence in human experiments seems to imply these are inhibitory processes (Sacchet et al., 2015a; Shin et al., 2017). Furthermore, the probability of the occurrence of a burst follows the trial-averaged beta power in the sensorimotor cortex; a rarefication of bursting events occurs during the ERD period and maximal likelihood coincides with the post movement beta rebound (Little et al., 2018). Furthermore, there are some early indications that these beta events form long-range transient synchronisations across the brain (Sacchet et al., 2015b), which could make bursts a potential mechanism of transient functional networks in the brain (O’Neill et al., 2018). There is also evidence that this bursting behaviour may facilitate communication to other regions of the body, as bursts centred around 20 Hz have been shown to be coherent with EMG recordings in the hand during a grip force task (Bourguignon et al., 2017). To be able to fully understand the mechanisms of these bursts, we require experimental validations for predictions which mathematical models of this bursting mechanism (Jones et al., 2009) make. Given the precise nature of single-unit INMS, it is a suitable experimental candidate to explore such mechanisms.

In a previous study by Kelly and colleagues (1997), INMS was combined with EEG, where they identified sustained evoked potentials that followed modulations in stimulation frequency. They found that both single-unit INMS and vibrotactile stimulation could evoke clear frequency following responses (FFRs) and these were matched to the stimulation frequency (33 Hz; Kelly et al., 1997). In our current study, we did not observe any such FFRs for neither INMS nor vibrotactile stimulation, although we used a stimulus frequency of 60 Hz. We can identify two potential reasons why this was the case, which are both linked to the experimental design. The first can be attributed to the number of trials per experiment. In the Kelly et al. study they used 200 trials of 1 s stimulation, whereas here we used 80 trials. Our choice of 80 was to optimise the experiment for capturing the modulation in beta power, which in the sensorimotor system takes several seconds to return to baseline after an event (Barratt et al., 2017; Fry et al., 2016; Pakenham et al., 2018; Robson et al., 2016), and so to gain an adequate inter-trial interval, the total number of trials is fewer. Assuming zero trials rejected, the Kelly et al. study has a 58% signal to noise ratio (SNR) improvement over our paradigm and this may be crucial to see such effects. The second reason, which may be more important than number of trials, is the stimulation frequency. Our choice of 60 Hz stimulation was driven by this providing a clearly perceivable, constant sensation for subjects when stimulating both FA1 and SA1 afferents (Johnson, 2001). However, it is likely that this frequency of stimulation is too high to generate a (detectible) FFR within the somatosensory cortex with MEG. Previous literature on the non-invasive imaging mechanoreceptive FFRs (see Vlaar et al., 2015 for a brief review) implies that the bulk of studies have been performed at a stimulation frequency of less than 40 Hz, and those which use higher carrier frequencies (such as Tobimatsu et al., 1999) used amplitude modulation below 40 Hz. Studies characterising the optimal frequencies show that the optimal frequency for an FFR is around 20-26 Hz (Jamali and Ross, 2013; Snyder, 1992; Tobimatsu et al., 1999). There has been a single report into the detection of subharmonics (Langdon et al., 2011), but their carrier frequencies were in this optimal spectral window.

In the present study, we did not perform a systematic analysis of the early somatosensory evoked fields since the experimental paradigm was designed to assess oscillatory responses. Ideally, single pulses would be used in an evoked fields analysis. In single pulse INMS, it is difficult for the subject to focus on such a brief sensation (i.e. to detect a single pulse coming from a single afferent), and such sensations have only been reported when stimulating FA1 units (Vallbo et al., 1984b). In this first study using MEG to study cortical responses to INMS, we wanted to ensure that we could gain a perceptual response to test, thus we delivered a longer train of pulses known to elicit perceivable sensations and detectable brain responses in EEG (Kelly et al., 1997). The main disadvantage of using such a train of pulses is that evoked fields from consecutive stimulation pulses will superimpose. Given that in our study we only had ∼16.6 ms between pulses, it would be difficult to deconvolve and resolve specific waveforms (e.g. M20, M50). To specifically investigate evoked fields using trains of stimuli, ∼500 ms intervals between pulses is required and will be of interest to address in future work. Nevertheless, we acknowledge whether single unit INMS can generate detectable evoked fields will be an interest to many, and have therefore performed a basic analysis of early evoked responses (see Appendix). Even with our suboptimal paradigm, we were able to localise the strongest source activations in the first 100 ms to S1 and S2. Therefore, this provides the opportunity of investigating whether differences between classes of mechanoreceptors are encoded in the earliest neural responses, or in later and slower oscillatory power modulations.

In our current study, we used bespoke flexible headcasts to improve the reliability of the spatial localisation of the oscillatory sources and comparison of INMS and vibrotactile responses between MEG scan sessions. The use of such headcasts has been shown to minimise head movement and the inter-session errors which arise from sensor-to-anatomy co-registration (Liuzzi et al., 2017). Since it was unknown how spatiotemporal responses to single-unit INMS would manifest in the human cortex, we wanted to ensure that we maximized the robustness of localisation of the sources between INMS and vibrotactile sessions using optimal methods. However, given that one unit (Unit 2a) was captured without a headcast and was localised to somatosensory cortex (see Supplementary material), with a source amplitude similar to that of the vibrotactile stimulation, future studies may not necessarily require a headcast, as long as sufficient practices in reducing head motion and co-registration errors are adhered to. Further the use of the headcast required care when inserting the participant in into the MEG sensor array once a unit was found, which has the potential of causing small electrode movements, which may lead to the loss of the single-unit nature of the stimulation. We originally intended to perform the search for mechanoreceptive afferents whilst the participant was within the MEG sensor array, however due to an unacceptable high level of capacitively-coupled mains-borne interference related to the MEG system earth in our installation this interference obscured the neural responses, making effective identification of single-unit recordings near impossible whilst the head was inside the MEG sensor array. The reference and recording electrodes used for INMS have very different impedances which makes the Common Mode Rejection Ratio performance for the pre-amplifier quite poor, but usually in screened room environments this is not an issue. We ameliorated this problem by lowering the participant out of the MEG sensor array and cutting the power to the MEG gantry during the search, with the former of these solutions necessitating careful entry and exit into the sensor array before and after data collection. It should be noted that cryogen-free on-scalp MEG solutions are starting to come into use (Boto et al., 2018, 2017), and since they provide of the order of 5-8 fold improvement in SNR over the central gyri and have sensors which move with the head, these will potentially provide flexibility for combined single-unit INMS without compromising on spatial specificity (Boto et al., 2016; Iivanainen et al., 2017).

In conclusion, combining the spatiotemporal specificity of MEG with the stimulation specificity of single-unit INMS enables the interrogation of the central representations of different aspects of tactile afferent signalling within the human cortices. Our initial findings imply that beta power in the somatosensory cortex is significantly modulated from the stimulation of single mechanoreceptive afferents, and given the precise nature of the stimulation method, single-unit INMS may serve as a controlled means to explore the mechanisms behind beta bursting events.

## Supporting information

## Acknowledgements

This work was supported by the Medical Research Council [grant number MR/M022722/1, to STF], the Swedish Research Council [grant 2017-01717, to JW], and the Knut and Alice Wallenberg Foundation [project NeuroSQUID, to JW] We also acknowledge the Medical Research Council MEG UK partnership [grant number MR/K005464/1]. Finally, the authors would also like to thank Kevin Aquino for his guidance on some of the early Freesurfer work and the online interactive plots.

## Appendix: A preliminary analysis into the early onset responses of INMS

As stated within the main manuscript, our experimental design means that our study is not particularly optimised for characterising the early onset somatosensory evoked fields; with only ∼16.6 ms between stimulation pulses, it would be difficult to deconvolve the effects of the adjacent stimulations. This will be addressed in future study, but for the time being we present a preliminary investigation of the early onset responses from our 10 FA1 units.

### Methods

In this analysis we present the evoked response to the entire train of stimulation pulses, rather than individual stimulations. Our methods to analyse the data are in places different to the analysis in the main manuscript and so a description follows. After the removal of artefacts from the data, we filtered between 1-80 Hz. For source reconstruction, we used the same source locations on the cortical surface and same forward model as before (3-shell BEM), but instead opted to use minimum norm estimation rather than a beamformer for inverse modelling as they are better suited to localising evoked sources (Ou et al., 2009). In particular, we used the sLORETA variant (Pascual-Marqui, 2002), where the sensor level noise covariance matric was generated from data 2 seconds prior to the stimulation onset. For a given vertex location *r*, three sets of source reconstruction weights representing three orthogonal dipoles were generated, which we shall define as *w*_*rx*_, *w*_*ry*_ and *w*_*rz*_ respectively. The sLORETA score is then calculated for given time point *t* by

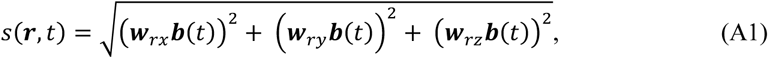

where *b* is the MEG sensor level data.

### Results

The results of this analysis can be found in Figure A1. Before focusing on the source level results, Figure A1A represents grand average sensor level plot, across all trials and subjects. We see there is a clear modulation of power oscillatory power within the 0-1 s epoch of microsimulation, both reflected in the higher amplitudes in the butterfly plot (coloured traces) and the increase in global field power (black trace) before eventually recovering back to baseline levels. Figure A1B is the source localised image of the earliest evoked responses in the contralateral hemisphere, averaged across both subjects and the first 100 ms of stimulation, with a threshold of 80% of the maximum sLORETA value applied. We observed two anatomical regions where activation is the strongest, the postcentral gyrus (S1) and the supramarginal gyrus (S2). Overlaid on top of the source plot in black is the fMRI derived hand region, which shows many of the peaks from the source plot contained within. The finding that the hand area intersected with this early source activation map and our areas of maximal beta power reduction (in Figures 3 and 4) increasing our confidence that those results were due to the stimulation and not other errant effects. Figure A1C and A1D show the average sLORETA timecourses in S1 and S2 in both hemispheres. We found that there were two distinct events in the data, a first early component that peaked in intensity at ∼200 ms, and a broader latent component towards the end of the stimulation. Note that the first component is also present in the main manuscript, represented as am increase in oscillatory power below 10 Hz (see Figure 3E). We also saw a bilateral response, with diminished responses in the ipsilateral (left) S1 and S2, which also appear to be lagged in time. This bilateral S2 response has been observed in somatosensory stimulation before (Gao et al., 2015). The latent effect of the ipsilateral hemisphere can be seen clearer in Figure A1D. From these results we are confident that with careful experimental design, evoked components related to single unit stimulation have the potential to be detected and characterised.

**Figure A1.**
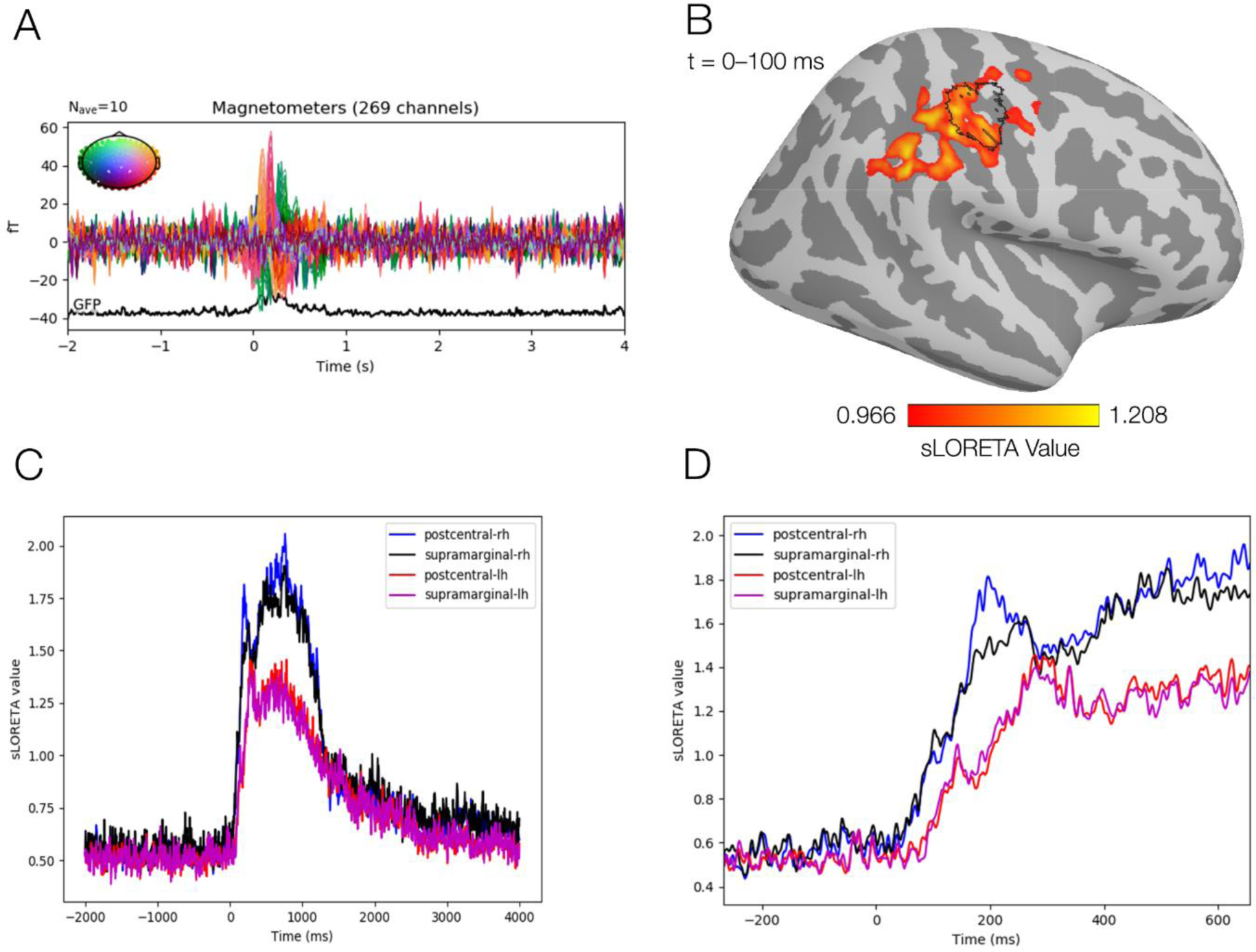
Results of a preliminary analysis into the early onset responses from INMS. A) A sensor-level grand average of the 10 FA1 units. B) A grand average sensor level plot of the first 100 ms of stimulation, showing the strongest sources in contralateral S1 (postcentral gyrus) and S2 (supramarginal) regions. For reference, the black overlay is the fMRI derived hand area (Sengupta et al., 2018). C) The average sLORETA timecourses from the left and right S1 and S2 areas. D) The same timecourses zoomed in to the first 600 ms, which shows the delayed ipsilateral response from both S1 and S2.

