## Supplementary material for "Imaging human cortical responses to intraneural microstimulation using magnetoencephalography"

George C. O'Neill[a*], Johan Wessberg[b], Roger H. Watkins[b], Rochelle Ackerley[bc], Eleanor L. Barratt[a], Paul M. Glover[a], Ayan Sengupta[a], Matthew J. Brookes[a], Rosa Maria Sanchez Panchuelo[a], Susan T. Francis[a]

[a]*Sir Peter Mansfield Imaging Centre, School of Physics and Astronomy, University of Nottingham, Nottingham, UK*
[b]*Department of Physiology, University of Gothenburg, Gothenburg, Sweden*
[c]*Laboratoire de Neurosciences Integratives et Adaptatives, Aix-Marseille Universite - CNRS, Marseille, France*

---

**1. Localising significant power changes in other epochs and frequency bands.**

Figures S1 and S2 show the results of applying our analysis pipeline and statistical tests in three other conditions for both INMS and vibrotactile stimulation. The conditions are as follows: alpha band power during stimulation, alpha power 1-2 seconds *after* stimulation and beta power 1-2 seconds *after* stimulation. In the alpha band we see that there is a group level significant reduction of power in the bilateral somatosensory during stimulation, but individually there is maximal overlap of 3/10 units, suggesting that this is a large effect in only a small subset of units than rather a universal feature. In post stimulation alpha we see nothing of significance in the sensorimotor areas. In post stimulation beta we see a significant increase power at the group level, which is localised to the contralateral somatosensory and motor cortices, but again, this is only significant in a small subset of units, all of which are were from the same subject (see figure S4 for an individual breakdown of unit activation locations).

We observe similar results when assessing the same band and epochs in the vibrotactile data (Figure S2), we again see event related desynchonisation of alpha power during stimulation at the group level, but only 2 units show individual significance in the somatosensory cortices. Post stimulation alpha shows (unlike in INMS) significant group level resynchronisation but no single unit shows significance in that region. Our post stimulation beta (much like the in the INMS) shows a group level significance, but only 3 units overlap.

**2. Localisation of individual units.**

Figures S3 and S4 are individual activation maps of INMS on the 10 FA1 units across two epochs in the beta band, during stimulation and 1-2 seconds *after* stimulation. All maps have a statistical threshold of $p_{corrected} < 0.05$. In Figure S3, we see that of the 10 units, 8 show any form of significant reduction in beta power during stimulation. Overlaid in red is a probabilistic atlas of the digits of the left hand (Sengupta et al., 2018), with the boundaries set to a 50% probability of being located within the digit area. All 8 units which show any form of significance intersect with this region.

Figure S4 shows the beta power from the 10 FA1 units in the resynchronisation period. Here the digit area instead has a black outline to not clash with the red of the statistical maps. Here we see that only three of the units show significant increases in power, all three coming from subject 4.

Figure S5 shows the significant beta power changes for the non-FA1 units.

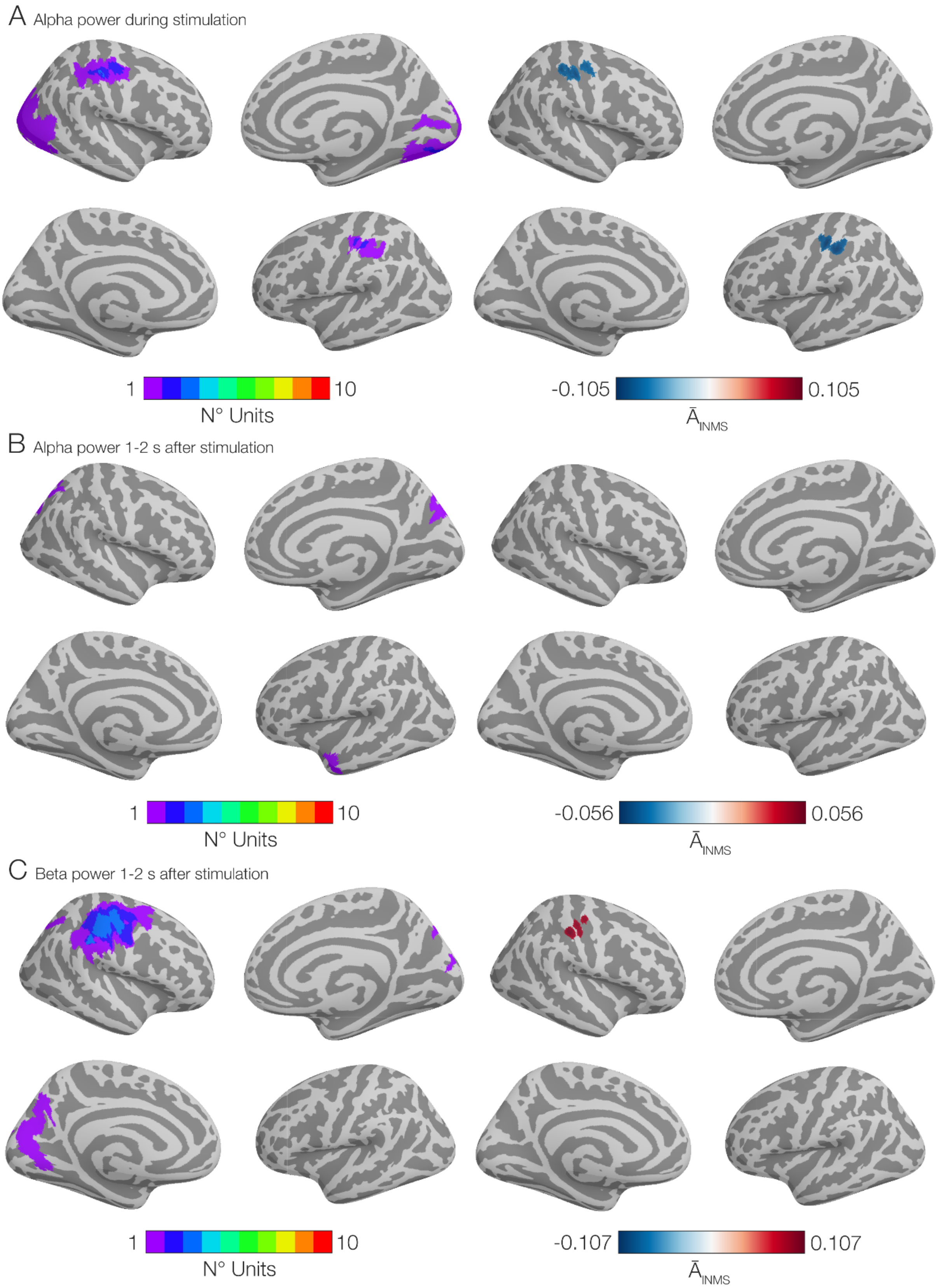


*Figure S1: Results of INMS source localisation in other epochs.*

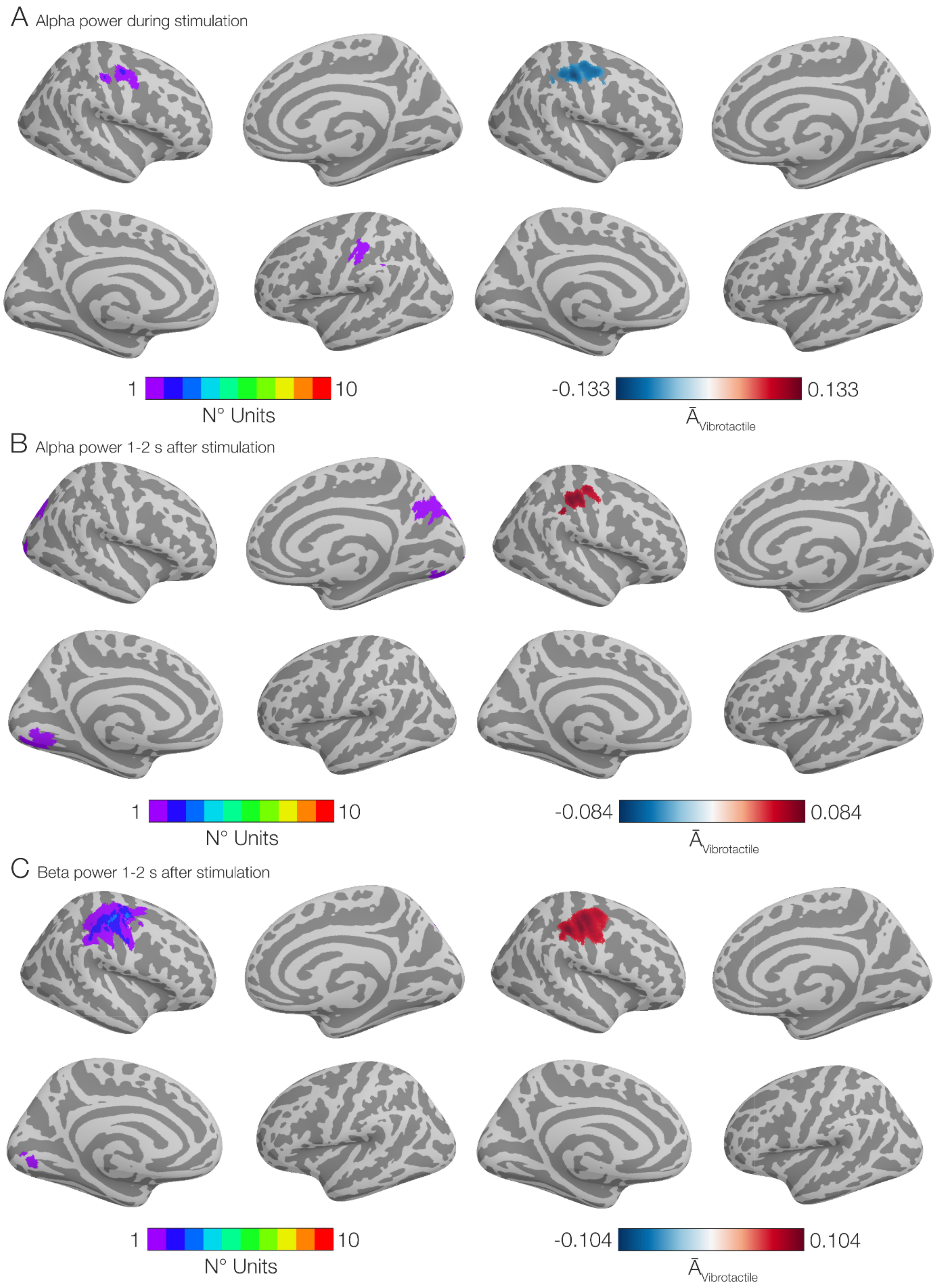


*Figure S2: Results of vibrotactile source localisation in other epochs.*

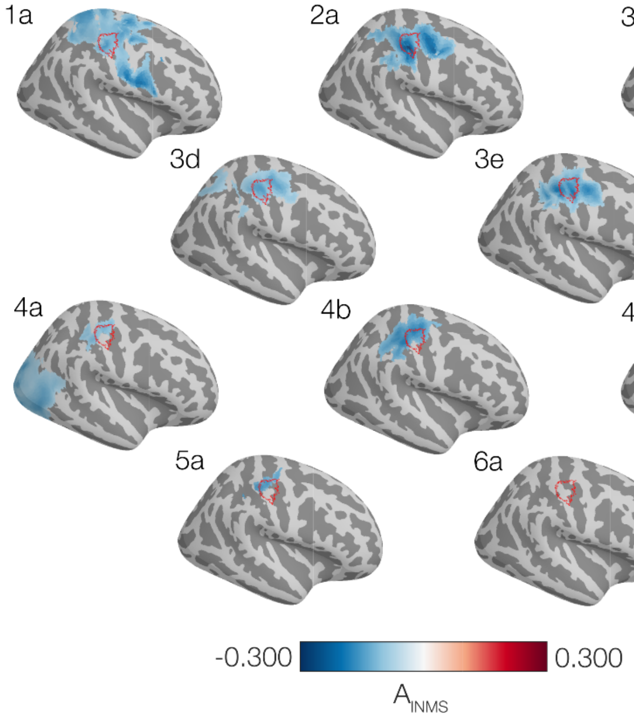


*Figure S3: Individual plots of the changes in beta power during INMS of the 10 FA1 units. Maps are threshold for significance.*

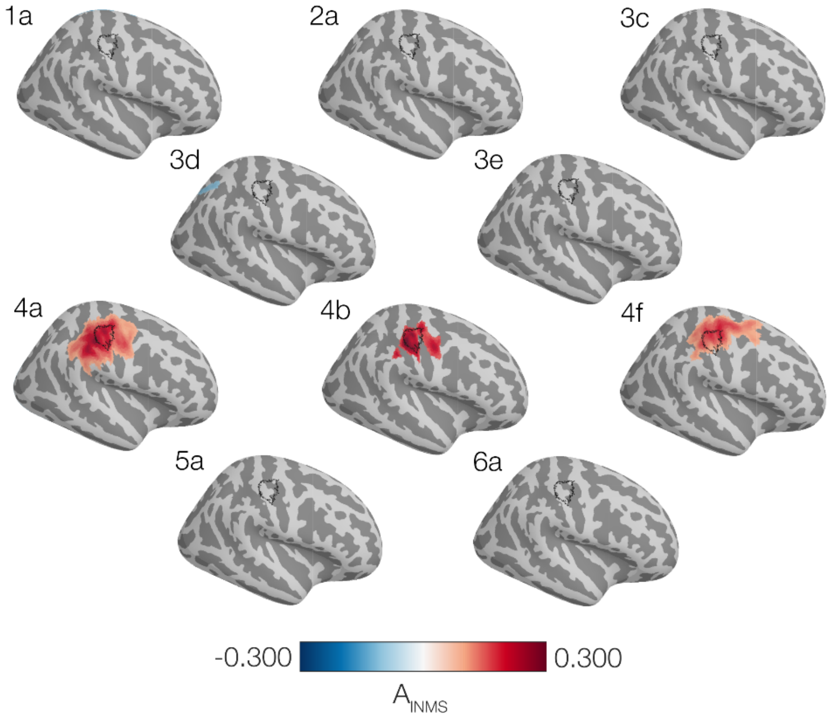


*Figure S4: Individual plots of the changes in beta power 1-2 seconds after INMS of the 10 FA1 units. Maps are threshold for significance.*

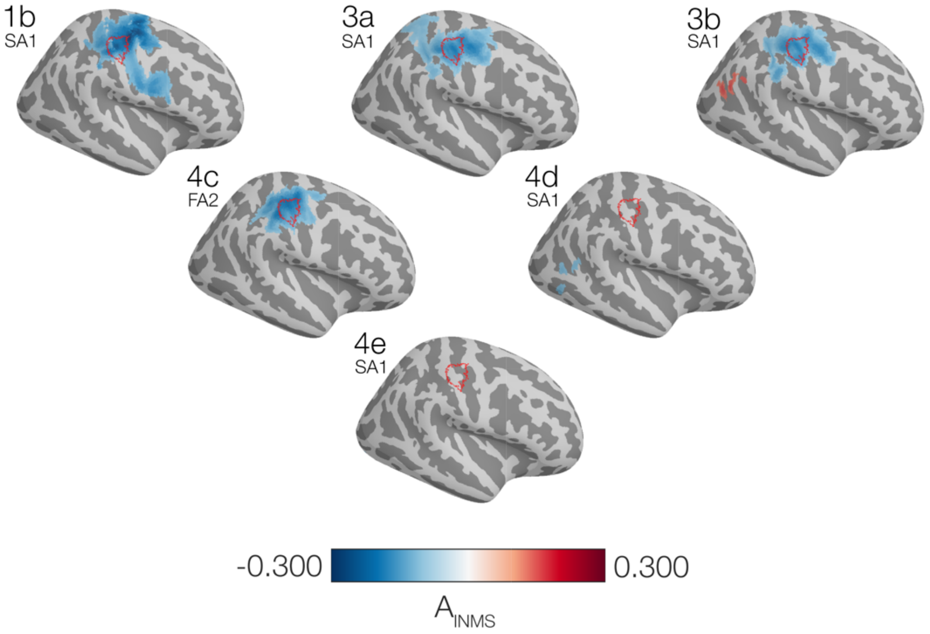


*Figure S5: Individual plots of the changes in beta power 1-2 seconds after INMS of the 6 non-FA1 units. Maps are threshold for significance.*
